# Second-generation transfer mediates efficient propagation of ICE*Bs1* in biofilms

**DOI:** 10.1101/2022.05.16.492222

**Authors:** Jean-Sébastien Bourassa, Gabriel Jeannotte, Sandrine Lebel-Beaucage, Pascale B. Beauregard

## Abstract

Horizontal gene transfer (HGT) by integrative and conjugative elements (ICEs) is an important mechanism in the spread of antibiotic resistance genes. However, little is known about the spatiotemporal dynamic of ICEs propagation in bacterial biofilms, which are multicellular structures ubiquitous in the natural and clinical environment. Using a fluorescently marked ICE*Bs1*, we report here that its propagation in biofilms is favored in recipient cells expressing the biofilm matrix. Also, conjugation appears restricted to clusters of bacteria in a close neighborhood in which a high level of ICE*Bs1* transfer occurs. These conjugative clusters are heterogenously distributed in the biofilm, forming close to the air-biofilm interface. Importantly, we established that transconjugant cells are the main contributors to ICE*Bs1* propagation in biofilms. Our findings provide a novel spatiotemporal understanding of ICEs propagation in biofilms, which should have an important role in our understanding of horizontal gene transfer in relevant settings.

**Importance:** The transfer of mobile genetic elements between bacteria is the main cause of the spread of antibiotic resistance genes. While biofilms are the predominant bacterial lifestyle both in the environment and in clinical settings, their impact on the propagation of mobile genetic elements is still poorly understood. In this study, we examined the spatiotemporal propagation of the well-characterized integrative and conjugative element (ICE) ICE*Bs1*. Using the Gram-positive *B. subtilis*, we observed that the main actors of ICE*Bs1* propagation in biofilms are the newly formed transconjugants that allow rapid transfer of ICE*Bs1* to new recipients. Our study provides a better understanding of the spatiotemporal dynamic of conjugative transfer in biofilms.

## Introduction

Horizontal gene transfer (HGT) is a fundamental phenomenon that drives the adaptation and evolution of bacteria in their environment (1). Conjugation is a preeminent HGT mechanism (2) that mediates the transfer of genetic material from a donor to a recipient cell upon direct cellular contact (3, 4). Since they encode complete mating machinery, conjugative plasmids and integrative and conjugative elements (ICEs) are genetic elements capable of self-transfer. They often carry accessory genes involved in metabolism, antibiotic resistance, and/or pathogenicity, which confer the cells bearing them a selective advantage (5–7). Consequently, they represent important actors in the emergence of multidrug resistance bacteria (8).

ICE*Bs1* is a 20.5 kb ICE present in multiple strains of *Bacillus subtilis*, a low G+C Gram-positive bacterium also able to form robust biofilms (9, 10). Since ICE*Bs1* gene function and regulation very well understood, it constitutes an excellent model to study the propagation of ICE in Gram-positive bacteria (11). Activation of ICE*Bs1* can result from the activation of the SOS response following DNA damages, in a RecA-dependant fashion (9, 11, 12). ICE*Bs1* activation is also mediated by RapI, a protein from the tetratricopeptide-repeat family encoded on ICE*Bs1* whose activity is inhibited by the small quorum-sensing peptide PhrI encoded downstream of *rapI* (9). Thus, the extracellular concentration of PhrI increases with the number of ICE*Bs1*-containing cells in a population, ensuring that the transfer will not be activated if closeby cells already contain the element (9). Since PhrI is taken up prior to sensing, its propagation in space is limited which leads to short-range communication (13).

Upon activation, ICE*Bs1* is excised from its integration site downstream of *trnS-leu2*, circularized, and then replicated by a rolling circle mechanism initiated at its origin of transfer (*oriT*) by the relaxase NicK (14). NicK is also involved in single-strand cleavage of the *oriT* to initiate the transfer via the type IV secretion system (T4SS) encoded by ICE*Bs1*, and the single-strand DNA is translocated to the recipient bacteria (15, 16). ICE*Bs1* is then recircularized before the complementary strand is synthesized (17–19). The newly completed ICE*Bs1* then integrates the chromosome via the *attB* site in the recipient genome. Importantly, transconjugants can also be immediately involved in another transfer to neighboring bacteria (3). In addition to the RapI-PhrI signaling, two other mechanisms limit ICE*Bs1* transfer to cells already bearing a copy of the element (20). The repressor ImmR mediates an immunity analogous to phage immunity (12), while YddJ, also encoded on ICE*Bs1*, mediates an exclusion mechanism by inhibiting the transfer from the ICE*Bs1* conjugation machinery (20).

Bacterial biofilms are microbial communities surrounded by a self-produced extracellular matrix (21, 22). Biofilms are ubiquitous in the environment and are involved in most chronic bacterial infections (23). Bacteria within these multicellular structures have an increased tolerance to antimicrobials mainly due to the surrounding matrix, and this densely packed community provides rich intercellular interactions (22–25). Because of these characteristics, biofilms are favorable environments for HGT (4, 26, 27). Indeed, population-level analysis revealed that the transfer of conjugative plasmids was increased in various biofilms such as activated sludge communities, which are well-studied environmental biofilms (28–30). Examination of conjugative plasmids propagation in biofilms using fluorescence microscopy and microfluidic revealed a low infiltration of the plasmids in an already established biofilm (29). In some cases, this poor efficacy was attributed to the low metabolic activity of recipient cells (31). However, other studies suggested that a low nutrient availability did not affect the transfer ability (26). While the capacity of conjugative plasmids to invade a biofilm appears limited, analysis of the conjugative plasmids RP4 and pKJK5 displayed efficient transfer in growing biofilms (28, 32). Of note, transconjugant cells were shown to have an important role in driving the transfer of RP4 in dual-species biofilm while having a minor impact in a complex activated sludge community (28). While these observations led to a better understanding of conjugation in biofilms, the spatiotemporal dynamics and factors impacting the conjugative transfer of ICEs in these multicellular communities have not yet been characterized (27).

We previously demonstrated that *B. subtilis* biofilm formation promotes the transfer of ICE*Bs1* and that the production of the biofilm matrix exopolysaccharide (EPS) and amyloid-like fibers (TasA) is required for this high efficiency (33). This observation was corroborated by others who also showed that ICE*Bs1* encodes for a protein, DevI, able to influence biofilm formation (34). Here, we took advantage of the high propagation of ICE*Bs1* in *B. subtilis* biofilms to better understand the spatiotemporal parameters of conjugation in biofilms using fluorescent microscopy. We report that although biofilms are considered a favorable environment for conjugative transfer, ICE*Bs1* propagates in confined, relatively small areas that display a strong transfer level. We also observed that most conjugation events occur near the air-biofilm interface and that transconjugant bacteria are the key actors in the propagation of ICE*Bs1* in biofilms.

## Results

### Bacteria-producing matrix components do not preferentially acquire ICE*Bs1*

In a previous study, we observed that the production of biofilm matrix by recipient cells drives the strong conjugative transfer of ICE*Bs1* in biofilm. To examine if this conjugation efficacy is due to the matrix itself or to the general phenotypic state of the cell, we performed a conjugation assay with recipient cells deleted for *sinR*, the transcriptional repressor of the operons encoding for exopolysaccharides (*epsA-O*) and protein fibers (*tapA-sipW-tasA*) of the matrix (35). Donor and recipient cells were mixed at a 1 to 5 ratio, and conjugation efficiency was examined after 20 h of incubation on a biofilm-inducing (MSgg) medium (36). As seen in Figure 1A, deletion of *sinR* and thus overproduction of the extracellular matrix by all recipient cells lead to a significant increase in conjugative transfer. Deletion of one or both matrix operons in a Δ*sinR* background caused a decrease in transfer which suggests that the production of these extracellular compounds, and not the phenotypic state, favors conjugation. To confirm the high ICE*Bs1* transfer to matrix-secreting recipient cells, we imaged conjugation and matrix production in parallel. The gene encoding for the red fluorescent protein mKate2 was inserted in ICE*Bs1*, which allowed us to follow its propagation by microscopy. We noticed that ICE*Bs1*-*mKate2* had a slightly diminished transfer level when compared to ICE*Bs1* bearing the *kan* resistance gene used previously, but still display robust conjugation (Figure S1). To visualize recipient bacteria, we integrated at a genomic locus the gene encoding for CFP fluorescent protein. Finally, the transcriptional reporter P_*tapA*_-*yfp*, which transcribes the *yfp* gene when cells are producing components of the biofilm matrix (24), was inserted in ICE*Bs1*-*mkate2* donors and *cfp* recipients cells. After a 20 h mating period, we inverted biofilms on a coverglass to directly image conjugation and determine the proportion of recipients and transconjugants expressing matrix genes (Figure 1B). Since the biofilm matrix has a strong positive effect on transfer, we expected the proportion of transconjugant cells to express P_*tapA*_-*yfp* more often than recipient cells. However, we did not observe a significant difference in the proportion of transconjugants cells expressing P_*tapA*_-*yfp* in comparison to the proportion observed for all recipient cells (Figure 1C). This discrepancy might be due to the fact that biofilm formation can be inhibited in transconjugants following the acquisition of ICE*Bs1* via expression of DevI, as previously reported by Jones *et al*., thus limiting the expression of the fluorescent reporter in transconjugant cells (34).

**Figure 1.**
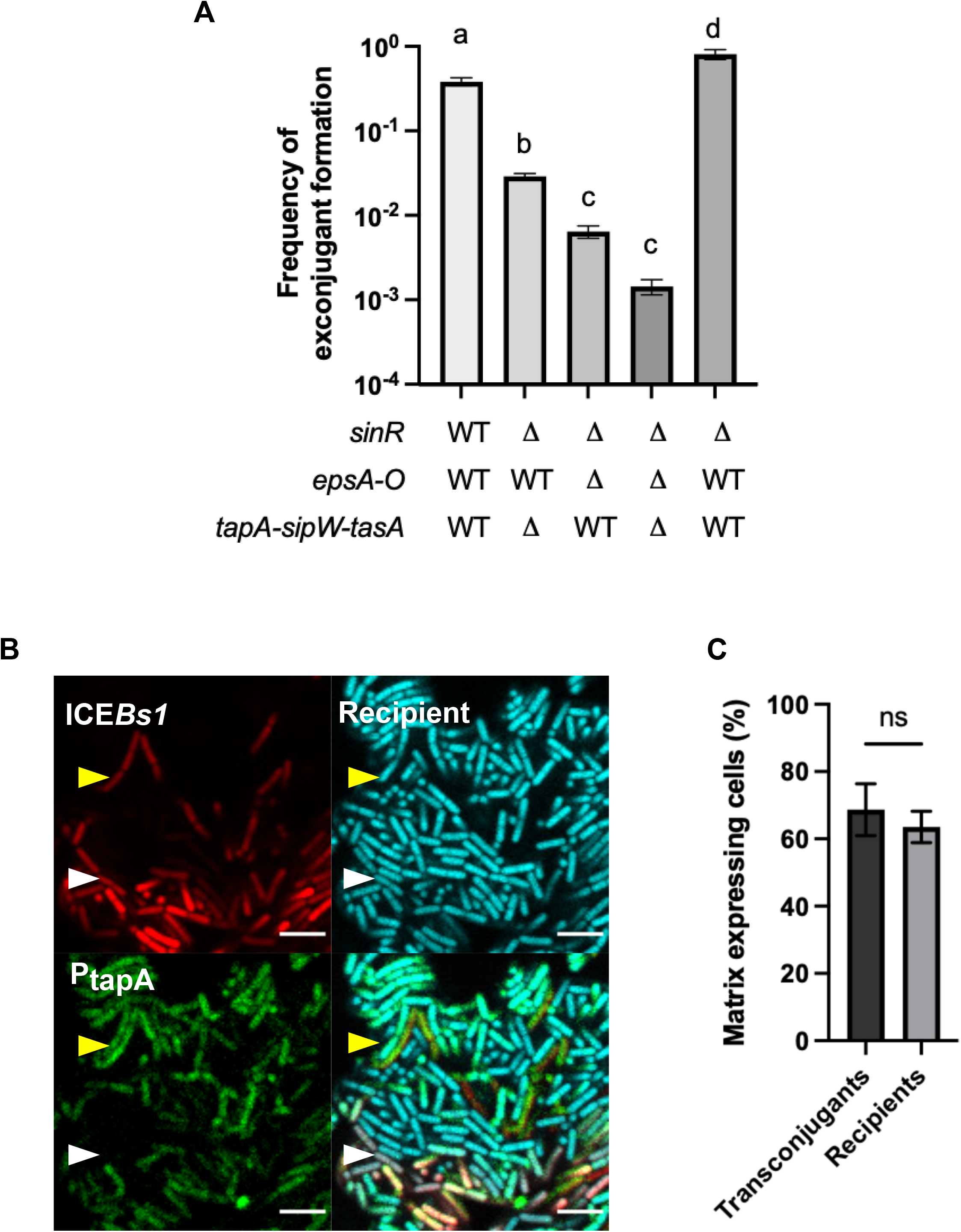
Influence of matrix production on ICE*Bs1* acquisition. (A) Mating between WT donor bacteria and recipient bacteria affected in their ability to produce biofilm matrix, incubated for 20 h on solid MSgg. The error bars represent the standard deviation of the mean (SD), and the results are representative of 3 biological replicates. Statistical analysis was done using a Brown-Forsythe and a Welch’s Anova followed by a Dunnett’s multiple comparaisons test with a statistical baseline of *P*≤0.05. (B) Mating between donor cells carrying ICE*Bs1*-*mkate2* (red) and recipient cells expressing *cfp* (blue) at a 1:5 donor to recipient ratio was incubated for 20 h on MSgg before imaging. Both donor and recipient also possess the P_*tapA*_*-yfp* (green) biofilm reporter. The scale bar indicates a size of 5 µm. The white arrow shows a transconjugant that does not produce matrix and the yellow arrow shows a transconjugant producing matrix. The image is representative of more than 20 pictures of conjugative clusters from 3 independent biological replicates. (C) The proportion of transconjugant and recipient cells expressing matrix gene, as determined by enumeration of 9 conjugative clusters using microscopy images. Statistical analysis was done using Paired *t*-test; ns *P*>0.05. Error bars represent the standard deviation (SD), and the results are representative of 3 independent replicates.

### ICE*Bs1* transfer in biofilm occurs in clusters

Intriguinly, microscopy observation revealed that conjugation events in biofilm appeared to be concentrated in regions of high transfer. To further examine this phenomenon, we used the ICE*Bs1*-*mKate2* reporter to image conjugation in cells incubated on biofilm and non-biofilm medium after 12 h, 16 h, and 20 h. In these assays, genes encoding for CFP or GFP fluorescent proteins were integrated at a genomic locus of donor and recipient bacteria, respectively. After 12 h of incubation, we observed that ICE*Bs1* transfer was initiated in a few donor bacteria with strong mKate2 fluorescence scattered through the biofilm, while most donor cells showed low fluorescent red signal (Figure 2A). This discrepancy between the fluorescence of various donor cells is likely due to the activation of ICE*Bs1* and its subsequent replication, which would increase the number of *mKate2* copies (37, 38). Four hours later, those initial donors had transferred ICE*Bs1* to neighboring cells, forming clusters of mKate2+ cells (Figure 2A). By 20 h, the clusters were enlarged and almost all the bacteria included in the area were transconjugant cells, many of which also strongly expressed mKate2 (Figure 2A). The propagation of ICE*Bs*1 was restricted to these active conjugative clusters (defined as containing at least 5 transconjugants) formed by donor cells surrounded by transconjugants.

**Figure 2.**
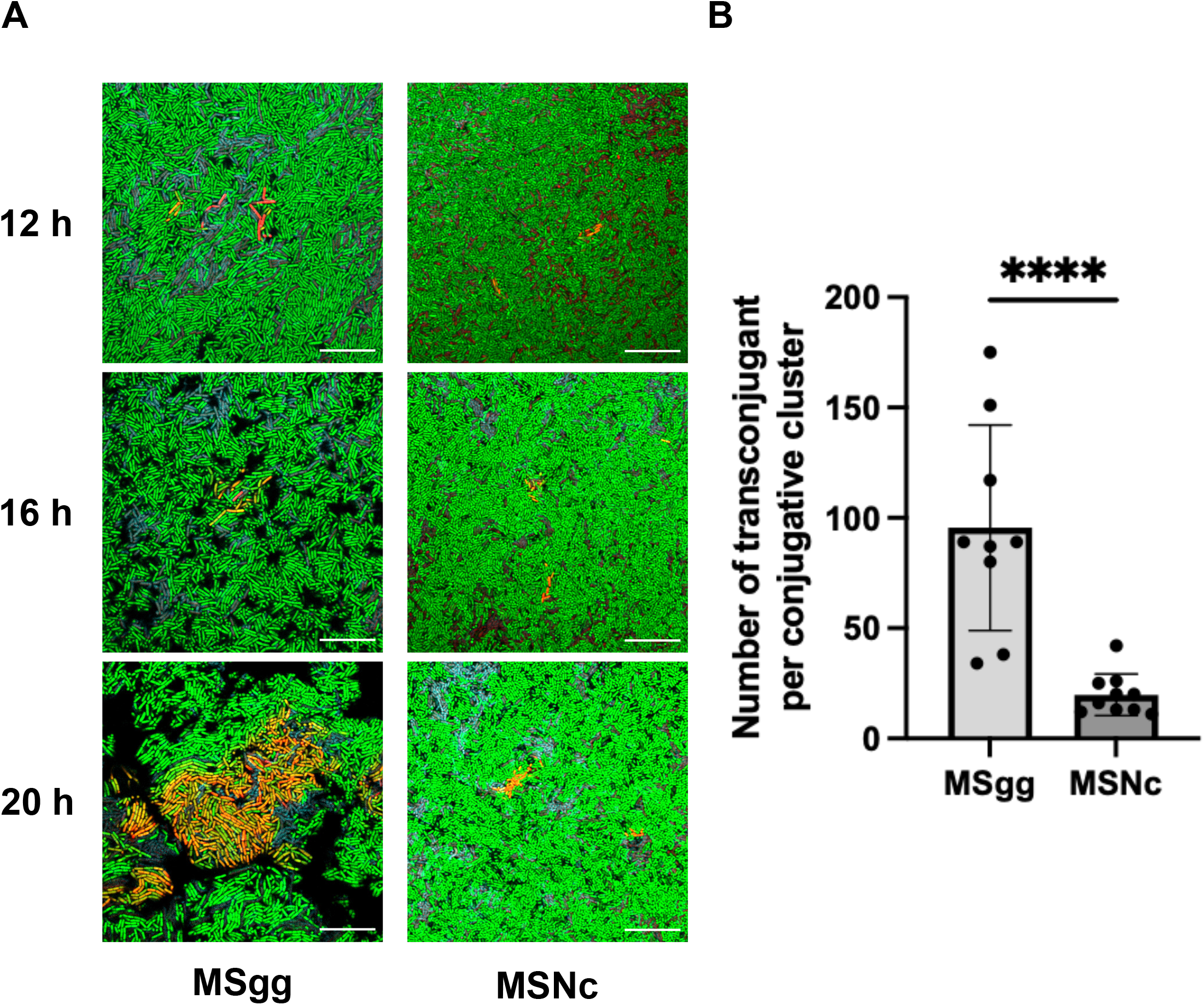
ICE*Bs1* transfer occurs in clusters. Donor strain expressing *cfp* from a genomic locus and bearing ICE*Bs1*-*mkate2* were mated at a 1:5 donor to recipient ratio with a recipient strain expressing *gfp* and visualized by fluorescent microscopy after 12 h, 16 h, and 20 h at 30 °C on biofilm (MSgg) and non-biofilm (MSNc) media. Donor cells appear blue to purple if the expression of mKate2 is low, and red if there is a strong mKate2 fluorescence; recipients are shown as green and transconjugants appear either orange if having a strong mKate2 fluorescence or yellow for a weak mKate2 fluorescence. Scale bars indicate a size of 10 µm. Pictures are representative of at least 9 pictures from 3 independent experiments. (B) Transconjugants composing the different conjugative clusters, as imaged in (A), were enumerated on biofilm (MSgg) and non-biofilm (MSNc) media. At least 9 pictures from 3 independent experiments were used, and each dot represents a cluster of at least 5 transconjugant cells. Statistical analysis was done using a *t-*test; **** *P<0*.*001*. Error bars represent the standard deviation (SD).

We previously showed that matings performed on a non-biofilm medium have a hundredfold less conjugation events than mating on a biofilm-inducing medium (33). Similar to the biofilm medium, at 12 h on a non-biofilm medium we observed few donor cells with a high level of mKate2 fluorescence and very few transconjugants. At 16 h and 20 h, ICE*Bs1* was also disseminated to neighboring cells but the conjugative clusters appeared significantly smaller than those formed on the biofilm medium (Figure 2A). Further analysis confirmed that at 20 h in biofilm conditions, the conjugative clusters were composed of an average of 95 cells while in non-biofilm conditions, they contained approximately 20 cells (Figure 2B). Our results show that ICE*Bs1* transfer in biofilm and non-biofilm conditions was confined into clusters, but these were significantly larger in biofilm conditions.

### ICE*Bs1* transfer in biofilm is heterogeneous

Imaging of biofilm inverted on a microscope coverslip does not recapitulate the complexity of the biofilm structure, which is a very heterogeneous environment. Thus, the presence of conjugative clusters in a fully formed colony biofilm was examined using transversal imaging. Matings were performed by dropping a mix of donor and recipient cells, bearing the same fluorescent reporters as described in Figure 2, at a 1 to 5 ratio on a solid biofilm-inducing medium. Biofilms were fixed and included in O.C.T. compound after 12 h, 16 h, 20 h, 24 h, and 28 h of incubation, and thin sections of the biofilms were prepared using a cryomicrotome to obtain slices containing a view of the entire depth of the biofilm. Slices were then mounted between a slide and the coverslip with an aqueous montage solution for imaging by confocal microscopy.

After 12 h of incubation, donor bacteria with strong mKate2 fluorescence were disseminated through the biofilm but there were few transconjugants, similar to what was observed in inverted biofilms. After 16 h of incubation, small independent clusters of mKate2+ (red) cells composed of donors and transconjugants were visible (Figure 3A, B). Longer biofilm incubation times led to larger clusters composed of transconjugant cells, suggesting a strong multiplication of transconjugants and/or a sustained transfer of ICE*Bs1*-*mkate2* (Figure 3A, B). Of note, most of these clusters appeared to have a vertical expansion and to be heterogeneously distributed in the biofilm. Positional analysis was performed on the clusters by measuring the distance between the top of the cluster and the air interface (Figure 3C; light blue), and between the bottom of the cluster and the medium interface (purple). The results clearly demonstrate that most clusters were found close to or directly at the air-biofilm interface of the biofilm, while the bottoms of the cluster were more randomly distributed respectively to the medium interface (Figure 3C).

**Figure 3.**
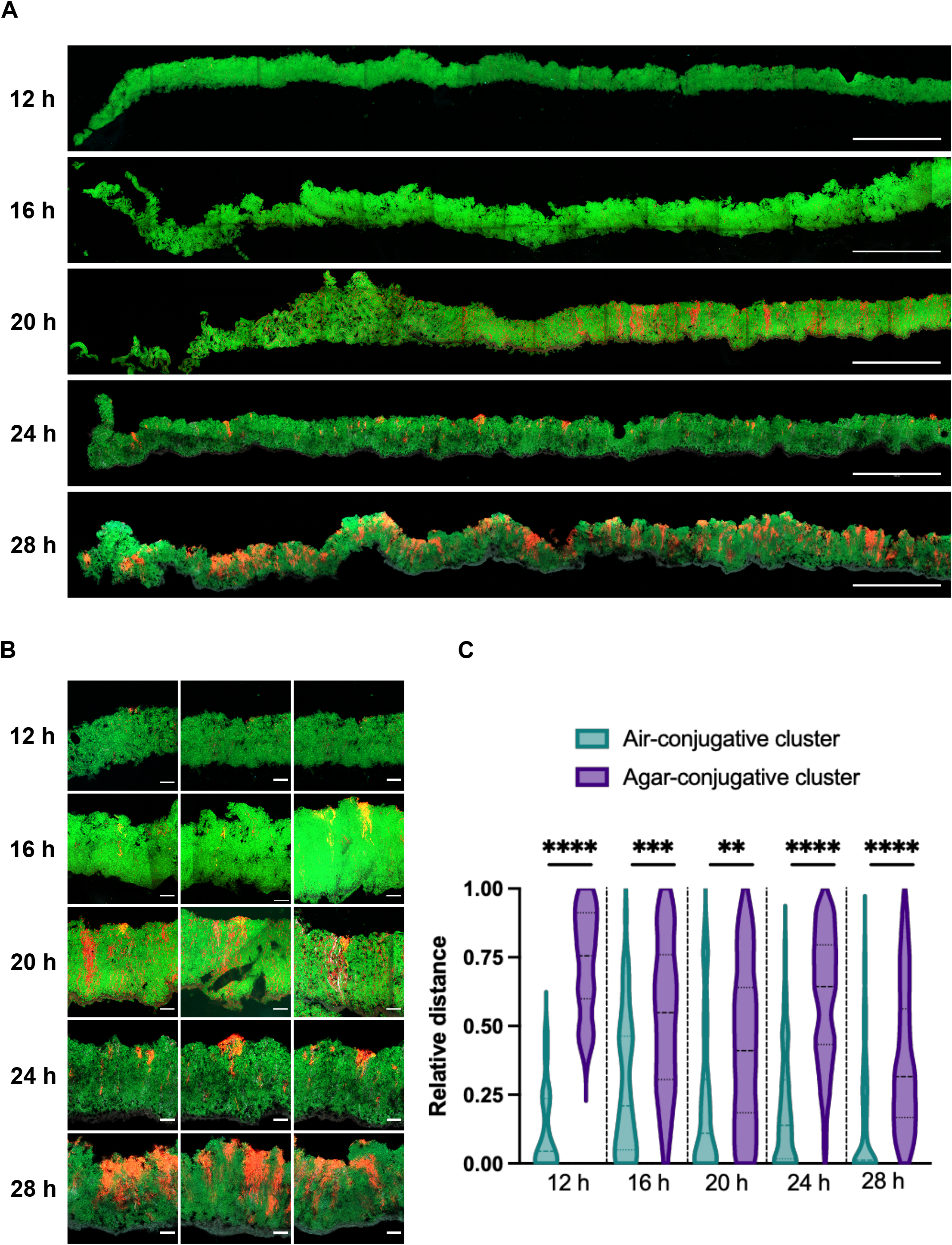
Spatiotemporal analysis of ICE*Bs1* propagation within a biofilm. Vertical thin section of biofilm composed of donor cells expressing ICE*Bs1*-*mkate2* with recipient cells expressing a *gfp* gene at a 1:5 donor to recipient ratio. At the indicated time, biofilms were fixed prior to cryosectioning. Wide view of the thin section representing approximately a third of a biofilm (A) and a close-up of the conjugative clusters (B). Donors appear red, recipients are shown in green and transconjugants are in yellow to orange color. The scale bars indicate a size of 250 µm (A) and 20 µm (B). (C) The relative distance of the conjugative cluster from the air-biofilm (light green) or agar-biofilm (purple) interface. Relative distance is reported as the distance separating the conjugative cluster to the interface, relative to the depth of the biofilm at the cluster’s location. The darker dotted lines represent the median and lighter dotted lines the 25^th^ and 75^th^ percentile. Results are a compilation of at least 19 conjugative clusters from at least 3 independent biofilms. Statistical analysis was done using a t-test, and shows that the bacterial cluster is closer to the air-biofilm interface than the agar-biofilm interface. (**, *P*<0.01; ***, *P*<0.001, ****, *P*<0.0001).

### Second-generation transfer mediates efficient ICE*Bs1* propagation inside the biofilm

Using 3D imaging of the conjugative cluster on inverted biofilm, we noticed that many transconjugant cells were not adjacent to donor cells (Video S1). This observation suggests that the transfer of ICE*Bs1* mediated by transconjugants could play an important role in ICE*Bs1* propagation in biofilms. To examine the importance of this second-generation transfer, we designed a conjugation assay in which only the donor cells, but not the transconjugants, could transfer ICE*Bs1*. We first performed an in-frame markerless deletion of *conG*, a gene present on ICE*Bs1* which encodes for an essential protein of its conjugative machinery (16). The *conG* complementation was constructed in *trans* under the control of the inducible promotor P_*hyperspank*_. As previously reported, deletion of *conG* completely abrogated ICE*Bs1* transfer but the presence of *conG* in *trans* in both the donor and recipient cells restored conjugation to wild-type levels (Figure 4A). Importantly, if the complementation was only present in the donor cells, ICE*Bs1*Δ*conG* was able to transfer from donor cells to recipient cells, but the newly formed transconjugants were unable to further propagate ICE*Bs1*Δ*conG*. Using this approach, we observed a significant decrease in the conjugation efficiency in biofilm and non-biofilm conditions when *conG* was complemented only in donor cells, compared to when it was complemented in both donor and recipient cells (Figure 4A). This result confirmed that transconjugant bacteria propagate ICE*Bs1* after its acquisition. Importantly, transfer by transconjugant cells represented about 99% of ICE*Bs1* transfer in biofilm conditions, while it represented only 43% of the transfer in non-biofilm conditions (Figure 4B). These results indicate that ICE*Bs1* propagation in biofilm is mediated by a rapid spread via transconjugant cells.

**Figure 4.**
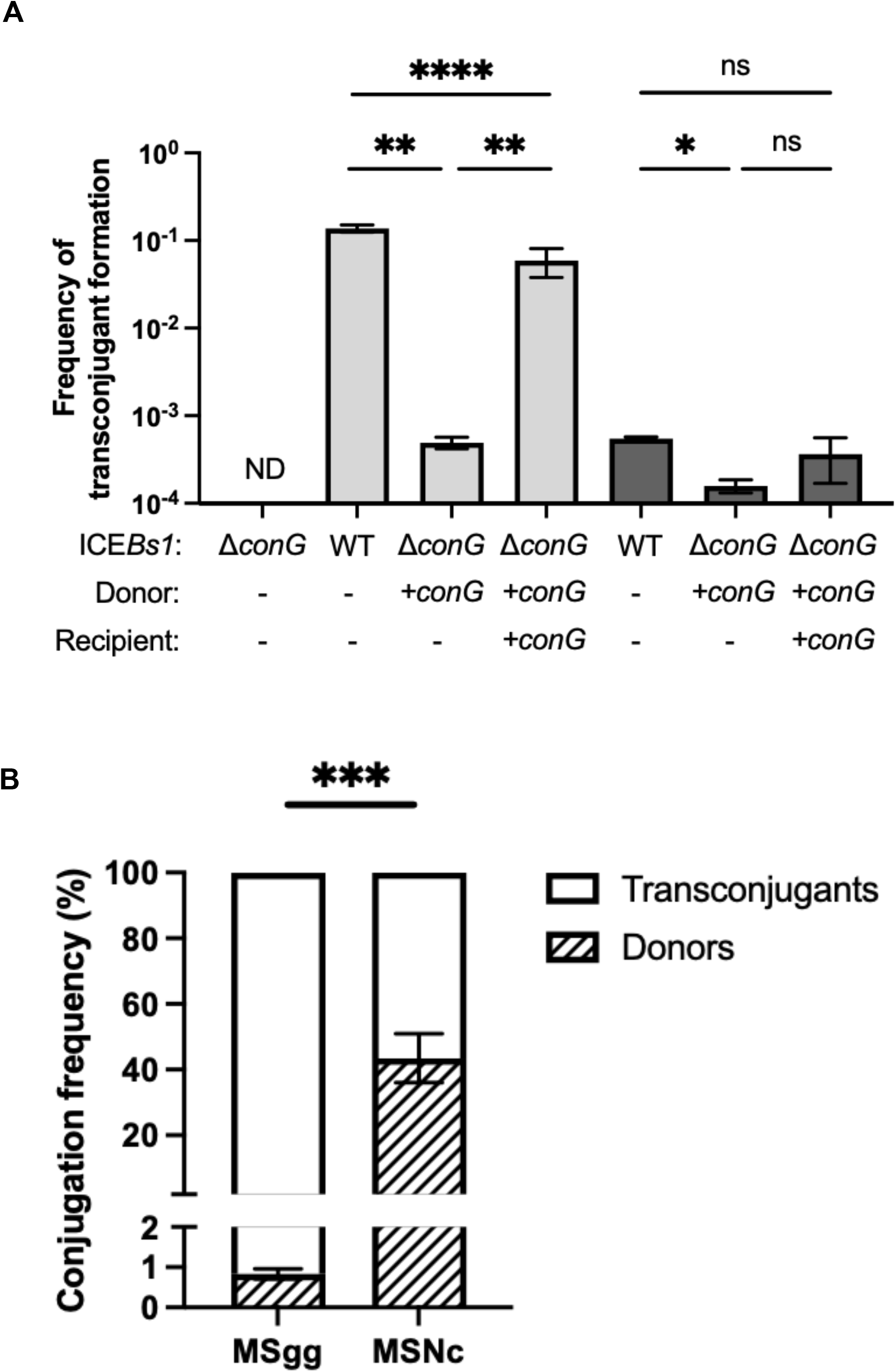
Transconjugants play a major role in ICE*Bs1* propagation. (A) Donor cells with an in-frame deletion of *conG* in ICEBs1 (Δ*conG*) were complemented *in trans* by expressing *conG* under the control of an IPTG-inducible promoter inserted at the *amyE* locus (+*conG*). Cells were mixed at a 1:5 donor to recipient ratio on a biofilm (MSgg, light gray) or a non-biofilm (MSNc, dark gray) media with 50 µM IPTG induction. Statistical analysis was performed using one-way ANOVA test followed by Tukey’s multiple comparison test ; *, *P*<0.05; **, *P*<0.01; ****, *P*<0.0001) (ND; non-detectable). (B) The proportion of conjugation attributed to donor and transconjugant cells was calculated by determining the relative contribution of donor conjugation (+*conG* in donor cells only) to total conjugation (+*conG* in donor and recipient cells). Statistical analysis was done using an unpaired *t*-test; *** *P<0*.*001*. Results showed are representative of at least three independent replicates.

It was previously reported that ICE*Bs1* can be efficiently transmitted via transconjugants by spreading rapidly through bacterial chain cells (3). Since *B. subtilis* can adopt this morphology in biofilm (36), we investigated if our observation of the importance of second-generation transfer could be explained by propagation in cell chains. Thus, we used mutants deleted for *lytC, lytD, lytE*, and *lytF*, which encode autolysins responsible for the peptidoglycan cleavage following cell division allowing the cells to separate (39, 40). While these deletion mutants forced the formation of cell chains (Figure S2A), they did not display an increase in ICE*Bs1* transfer in non-biofilm conditions (Figure 5A). Similarly, overexpression of the autolysin locus *lytABC* in biofilm formation conditions did not impair ICE*Bs1* ability to transfer in biofilms (Figure 5B) while reducing the presence of cell chains (Figure S2B). From these results, we conclude that ICE*Bs1* transfer by transconjugant is key for its efficient propagation in biofilms, but that this effect is not mediated by cell chains.

**Figure 5.**
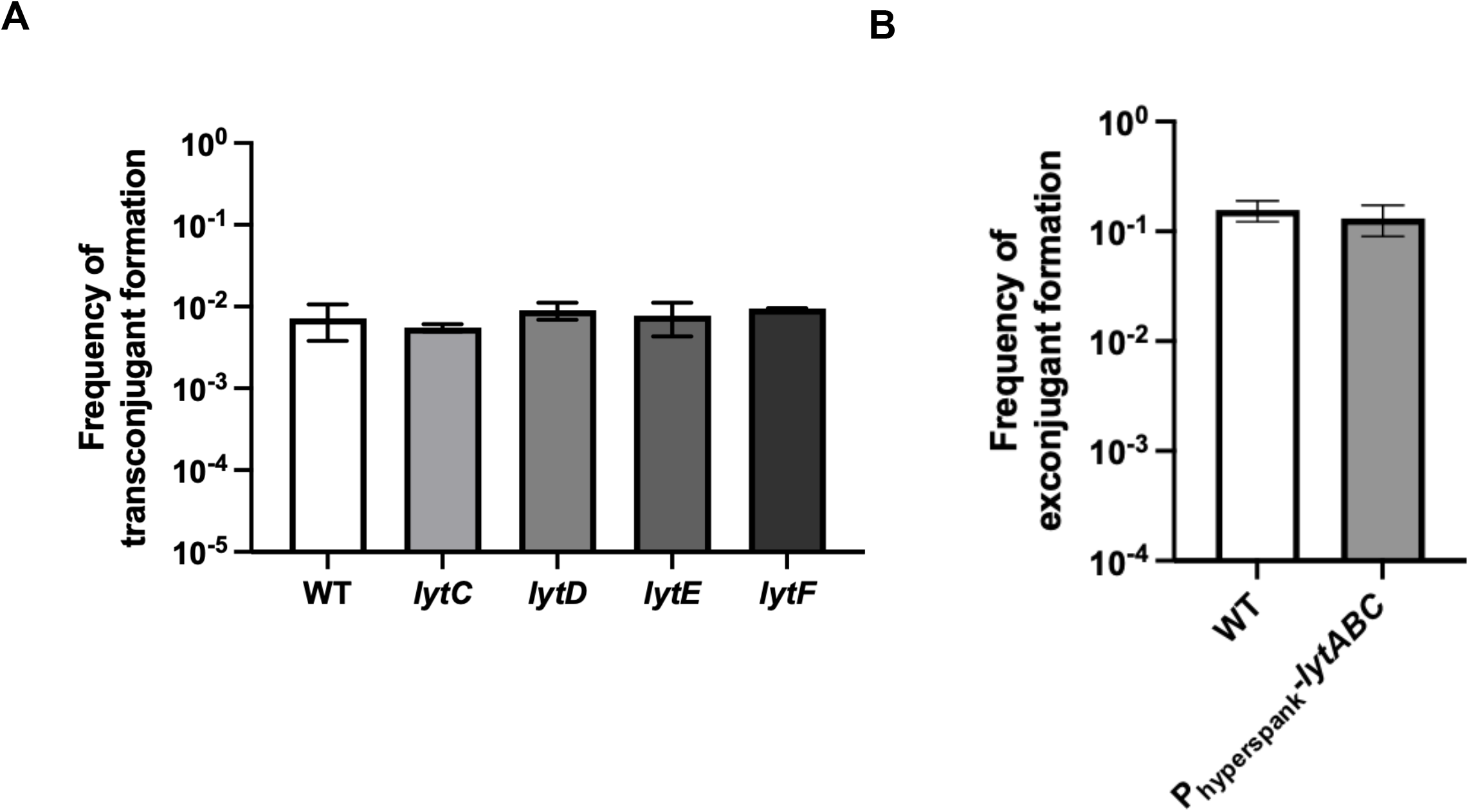
Cell chains are not involved in the propagation of ICE*Bs1* in a biofilm. (A) WT donor cells were mated with recipient cells lacking different autolysins and incubated on a non-biofilm (MSNc) medium. (B) WT donor cells were mated with recipient cells overexpressing *lytABC* under the control of an IPTG-inducible promoter and incubated on a biofilm-inducing (MSgg) medium with 50 µM IPTG. For (A) and (B) cells were mixed at a 1:5 donor to recipient ratio and incubated for 20 h at 30 °C. Statistical analysis was performed using an ANOVA for (A) and *t test’s* for (B); (*ns P>0*.*05*), and showed no diffeernce. The results shown are representative of at least 3 independent experiments and error bars represent the standard deviation (SD).

## Discussion

The molecular mechanisms underlying conjugation are well understood for many mobile genetic elements, but little is known about the spatiotemporal parameters of their propagation in more natural contexts (27). Here, we used the well-characterized ICE*Bs1* and its host, *B. subtilis*, to demonstrate that although propagation of ICE*Bs1* in biofilm is extremely efficient, its transfer is confined to clusters of cells where ICE*Bs1* replicates and rapidly spread through transconjugant bacteria.

We recapitulated our previous observations that the production of matrix by recipient cells plays an important role in the propagation of ICE*Bs1* in biofilm. Using a deletion of *sinR* to force production of biofilm matrix, we uncoupled general phenotypic differentiation from *epsA-O* and *tapA-sipW-tasA* expression, further confirming the importance of the matrix components (Figure 1A). However, microscopy observations revealed that transconjugants did not systematically express matrix genes, which might reflect the activity of the ICE*Bs1* encoded *devI* as a Spo0A inhibitor. Our current hypothesis is that matrix would be required only at the beginning of transfer to stabilize the contact between donor and recipient cells.

We observed that ICE*Bs1* forms conjugative clusters which are mostly located close to the air-biofilm interface, showing a clear heterogeneity in vertical distribution (Figure 3C). Many of these conjugative clusters also display an uneven appearance with a preference for vertical expansion which could suggest a polar growth of the transconjugant cells, a polar transfer, or both (Figure 3B). Similarly, propagation of the TOL plasmid pWWO in an already formed biofilm also appeared to be confined to the few upper layers of cells at the liquid medium interface of the biofilm (29). A lack of nutrients and oxygen inside the biofilm was proposed as an explanation for the confine transfer of pWWO at the liquid medium interface since the conjugative transfer requires a high energy cost (22). In our mating assay on solid biofilm-inducing medium, nutrient availability does not appear to influence ICE*Bs1* conjugative transfer since the cells present in the upper part of a colony biofilm have access to the least nutrients. Indeed, sporulation occurs rapidely in this specific biofilm layer (24). However, the air-biofilm interface was also shown to be the site of early TasA expression and faster accumulation of the biofilm matrix (24), which could explain the preferential localization of clusters close to this interface prior to occurrence deeper in the biofilm.

We observed that initially, in biofilm, only a small number of bacteria produce a bright mKate2 fluorescent signal resulting from ICE*Bs1* excision and its replication in plasmid form (Figure 2). This low number is in agreement with our previous observation that ICE*Bs1* is excised in less than 0.1% of the donors in both biofilm and non-biofilm conditions (33). These initial events might be driven by the local density of donor and recipient cells since a high density of donor cells will lead to a repressing extracellular concentration of PhrI (9, 13). A similar observation was reported for the conjugative plasmid RP4 whose propagation in a growing biofilm also appears to arise from distinct areas prior to the expansion through most of the biofilm (28). Since transconjugant clusters can expand to almost a hundred cells, repression by PhrI is unlikely to have an effect on second-generation transfer. We hypothesize that this phenomenon could be explained by the kinetic repression by PhrI which needs to be synthesized, exported, and then imported back to inhibit RapI, a process that might be slower than the conjugative transfer of an already active ICE*Bs1* present in transconjugants.

ICE*Bs1* propagation mediated by transconjugant cells was previously shown, but its importance for overall conjugative transfer was not evaluated (3). Our approach allowed us to precisely determine that the transfer initiated by transconjugant cells represents more than 99% of the total conjugative events observed in a biofilm (Figure 4B). Surprisingly, even if ICE*Bs1* can rapidly spread through bacterial cell chains (3), this ability does not appear to contribute significantly to conjugation in biofilms (Figure 5).

The high transfer efficiency by transconjugants might be explained by the initial absence of the negative regulator ImmR, which is degraded upon ICE*Bs1* activation (41). Following translocation, ICE*Bs1* in its plasmid form will be able to maintain its activated state, replicating and transferring efficiently as proposed in another study (3). A high replication before the conjugative transfer was also observed for ICE*clc*, an ICE present in *Pseudomonas putida* that induces a transfer competent (tc) differentiation in donor bacteria after activation of ICE*clc* genes (42). Activation of these genes led to a transient replication step prior to the conjugative transfer (43). Thus, bacteria in which ICE*clc* replicates are most likely to propagate the element. While ICE*Bs1* replication in the donor bacteria was shown to be unnecessary for the conjugative transfer of this element (44), a high copy number of ICE*Bs1* in transconjugants could favor the high secondary transfer while also supporting integration in the genome (44, 45). Our current model is that the strong ICE*Bs1* transfer in biofilm results from the combination of high ICE*Bs1* activity in newly formed transconjugant cells with the stabilizing effect of the extracellular matrix on cell-to-cell contacts between transconjugants and recipient cells.

While ICE*Bs1* propagation is driven by transconjugants, it is not the case for most conjugative elements. Previous reports showed complete recovery of the conjugative levels of multiple elements upon deletion of a protein from their T4SS machinery and complementation only within the donor cells (46–48). These elements are members of multiple incompatibility groups present in both Gram-positive and Gram-negative bacteria, which indicates that a high level of second-generation transfer might be key for a certain subset of conjugative elements. However, these complementation assays were done in non-biofilm conditions; since our study demonstrates that the epidemic transfer is important, particularly in biofilm, more investigation might reveal if it is a biofilm-specific feature of conjugation or not.

Our study provides new insights into the spatiotemporal dynamic of conjugative element propagation in biofilms. Since biofilms are ubiquitous in clinical environments where multi-resistant bacteria can emerge, a better understanding of ICEs propagation in these conditions is required to develop efficient strategies to prevent or decrease horizontal gene transfer. Further studies on the epidemic transfer of ICEs to better understand its real impact in natural environments are warranted.

## Materials and methods

### Strains and media

The strains used in this study were derived from the ancestor strain NCIB3610 (see Table S1). The bacterial growth media used is LB (Luria Bertani; 1% tryptone, 0.5% yeast extract, 0.5% NaCl) and the different media used for mating assays are MSNc (5 mM potassium phosphate buffer, pH 7, 0.1 M morpholinepropanesulfonic acid [MOPS], pH 7, 2 mM MgCl_2_, 50 µM MnCl_2_, 1 µM ZnCl_2_, 2 µM thiamine, 700 µM CaCl_2_, 0.2% w/v NH_4_Cl, 0.5% w/v cellobiose) (49), and MSgg (5 mM potassium phosphate buffer, pH 7, 0.1 M MOPS, pH 7, 0.025 mM FeCl_3_, 2 mM MgCl_2_, 50 µM MnCl_2_,1 µM ZnCl_2_, 2 µM thiamine, 700 µM CaCl_2_, 0.5% v/v glycerol, 0.5% w/v glutamate) solidified with 1.5% w/v agar. When needed, the following antibiotics were added to the media: MLS (1 µg ml^-1^ erythromycin, 25 µg ml^-1^ lincomycin), spectinomycin (100 µg ml^-1^), chloramphenicol (5 µg ml^-1^) and kanamycin (10 µg ml^-1^).

### Strains and plasmids construction

Most *B. subtilis* strains were made by transferring genetic constructions into NCIB3610 using SPP1-mediated generalized transduction (50). JMA384 (ICE*Bs1::kan*) was a gift from Alan D. Grossman’s lab (Massachusetts Institute of Technology, MA); BJM396 (*lytC*), BJM402 (*lytD*), BJM76 (*lytE*), and BJM104 (*lytF*) were gifts from David Rudner’s lab (Harvard Medical School, MA). pminiMAD2 (51) was a gift from Richard Losick’s lab (Harvard, MA). The *Escherichia coli* strain used for routine cloning was NEB 5α (New England Biolabs). The plasmids used in this article are listed below (Table S2). *tapA-sinR* deletion was created by long fragment homology PCR. The flanking regions were amplified using P746-P747 and P748-P749 respectively and the *erm* was amplified from pDG646. Plasmid constructions were done using pDR111 as the backbone plasmid unless indicated. *mkate2* was cloned into ICE*Bs1* by subsequent cloning of upstream and downstream regions of *yddM-yddN*. The upstream fragment was amplified with P342 and P343 primers and inserted at the BamHI restriction site downstream of *lacI*. The downstream fragment was amplified with primers P344 and P496 and was inserted while replacing the *amyE down* homology fragment with the AsisI and SacI restriction enzymes. P_*hyperspank*_*-mkate2* was amplified from PB396 (*amyE::P*_*hyperspank*_*-mkate2*) with primers P507 and P508 and inserted between the EcoRI and SphI restriction sites of pDR111. To complement the *conG* deletion, *conG* was amplified with primers P719 and P673 and inserted between the SalI and SphI restriction site downstream of P_*hyperspank*_ inducible promotor on pDR111. P_*hyperspank*_*-cfp* was obtained by amplifying *cfp* from pKM008 with primers P615 and P616 and cloned between HindIII and SphI restriction sites. HindIII is present in the amplification and not on primer P615. Overexpression of *lytABC* was done by amplification of *lytABC* with P647 and P648 and inserted between SalI and SphI restriction site. Constructions were transferred in *B. subtilis* 168 through natural competency, verified by PCR, and then transferred in NCIB3610.

### Markerless deletion

Markerless deletion of *conG* was done using pminiMAD2 (51). Upstream and downstream fragments were amplified with primers P668 and P669 and with primers P670 and P674, respectively. The two fragments were fused by long-fragment homology PCR and inserted into pminiMAD2 between BamH*I* and EcoRI restriction sites. Plasmid pJSB19 was inserted into MM294, a *recA*+ *E. coli*, to obtain concatemer. pJSB19 was then inserted into JMA384 using natural competency and then incubated overnight with selection at 40 °C. Multiple colonies were then incubated in LB broth for 3-4 hours at room temperature and diluted into fresh LB broth overnight at room temperature. Excision and curing of the plasmid were achieved by successive passages and growth at 37 °C. Cells were then plated on LB agar and colonies were streaked on LB with and without MLS. Colonies that lost antibiotic resistance were then PCR verified to confirm the deletion of *conG*. Sequencing of the deleted locus was also performed.

### Biofilm matings

Donor and recipient cells were grown from a single colony in 3 ml LB broth at 37 °C to late exponential phase (3-4 h), diluted at an OD_600_ of 1.5 in PBS 1X (137 mM NaCl, 2.7 mM KCl, 10 mM Na_2_HPO_4,_ 1.8 mM KH_2_PO_4_, pH 7.4) and mixed at a 1:5 donor to recipient ratio unless indicated otherwise. Cells were then centrifuged for 3 min at 5,000 rpm, and the pellet was resuspended in 1/10^th^ of the volume. 10 µl of the mix was dropped onto the appropriate medium and incubated for 20 h (unless time specified [Figure 2, 3]) at 30 °C. When necessary, expression of the different constructs was induced by adding IPTG (Isopropyl β-D-1-thiogalactopyranoside) to the medium. Fluorescent constructions were induced by adding 50 µM IPTG to both the liquid preculture and biofilm medium (Figure 1, 3, 4, Figure S2, Video S1). Complementation of *conG* was done by adding 50 µM IPTG to the biofilm-inducing medium (Figure 5).

To determine conjugation levels, colonies were collected from agar media with a folded tip and put in 1 ml PBS 1X, then sonicated at 30% amplitude 20 s twice with a 1:1 pulse. Cell suspensions were serially diluted and plated on LB with the appropriate antibiotic and allowed to grow until the next day. Donors, recipients, and transconjugants CFU were then counted. The frequency of transconjugants formation was expressed as a function of the number of recipients CFU (number of transconjugants devised by the number of recipients CFU) because transconjugants are an important actor in the propagation of ICE*Bs1* (Figure 4).

### Thin section

Preparation of the mating assay was done as previously described, but 2 µl instead of 10 µl of the mixture was dropped on MSgg medium with 1.5% agar poured into 6-well plates. At the appropriate time, biofilms were fixed using paraformaldehyde 4% for 10 min and washed twice with PBS. Samples of biofilm and agar were harvested from the plates and transferred into a homemade aluminum foil mold, overlaid with Tissue Plus O.C.T. compound clear (#4585, Thermo Fisher), and fast-frozen on dry-ice. The frozen samples were stored at -80°C, before being sent to Plateforme d’histologie de l’Université de Sherbrooke and sliced into 8-µm-thick using a cryomicrotome set to - 20 °C. Thin sections were placed on a positively charged slide (Globe Scientific inc. #1358W) and an aqueous mounting medium was added (0.5% N-propyl gallate, 50% glycerol, PBS) and covered with a number 1.5 coverslip. Slides were kept at 4 °C prior to imaging. Thin sections were analyzed using an Olympus FV3000 confocal microscope at 60x magnification (PlanApo objective 60X/NA1.4) with the following laser; 405 nm, 488 nm, and 561 nm. Laser intensity and sensor sensitivity were the same in all conditions. The distances separating the conjugative clusters with either the air-biofilm or biofilm-media interface were mesureed using the ImageJ software. The relative distance was calculated as the distance measure divided by the total thickness of the biofilm at the area of the picture taken.

### Inverted microscopy on biofilm

Preparation of the mating assay was done as previously described, but 2 µl instead of 10 µl of the mixture was dropped on MSgg or MSNc medium with 1.5% agar. At the appropriate time point, agar containing the biofilm was cut with a razor blade. Excess agar was removed carefully before inversion of the whole biofilm into a 35 mm petri dish containing a coverglass number 1.5 (P35G-1.5-14C, MatTek corporation). Pictures of the inner part of the biofilms were taken using an Olympus FV3000 confocal microscope at 60x magnification (PlanApo objective 60X/NA1.4) with the following laser; 405 nm, 488 nm, and 561 nm. Laser intensity and sensor sensitivity were the same for all conditions. For statistical analysis, bacteria were counted using ImageJ software with a cell counter plugin.

### Statistical analysis

Statistical analyses were performed using GraphPad Prism 9. Comparisons were done as specified in the Figure legend.

## Acknowledgments

We thank A. D. Grossman, D. Rudner, and R. Lobsick for their kind gift of strains, and members of the Beauregard, Burrus, Rodrigue, Côté, and Malouin laboratories for helpful discussions, and Alain Lavigueur for his comments on the manuscript. This work was supported by a Team Project Grant (2019-PR-253077) from the Fond de Recherche du Québec - Nature et Technologies (FRQ-NT) to P.B.B. and by Master Degree Fellowships; Bourse d’excellence VoiceAge from la Fondation de l’Université de Sherbrooke and (271990) from the Fond de Recherche du Québec - Nature et Technologie (FRQ-NT) to J-S.B.

---

**Figure S1. ICE*Bs1::mkate2* transfer at a slightly lower efficiency than ICE*Bs1::kan***. Donor cells carrying ICE*Bs1::kan* or ICE*Bs1-mkate2* were mated on MSgg medium at 30 °C for 20 h. The statistical analysis shows that ICE*Bs1-mkate2* transfer is slightly less efficient than the WT ICE*Bs1::kan* but remains high (*t-test; ** P<0*.*01*). Error bars represent the standard deviation (SD). The result is representative of at least three independent replicates.

**Figure S2. Lytic enzyme mutant phenotype**. (A) Lytic mutants were grown on solid MSNc medium containing 50 µM IPTG and incubated at 30 °C for 20 h prior to inverting and imaging using confocal microscopy. (B) Overexpression of *lytABC* was grown on solid MSgg medium containing 50 µM IPTG and incubated at 30 °C for 20 h prior to inverting and imaging using confocal microscopy. The scales bars represent a size of 10 µm. All strains expressed mKate2 except the *lytABC* overexpression which expressed YFP for the imaging.

**Video S1. Multiple transconjugant bacteria are not adjacent to donor cells**. Donor bacteria bearing ICE*Bs1-mkate2* and expressing a CFP fluorescent protein were mated with recipients expressing a GFP fluorescent protein. Pictures were taken a in Z-stack of 10 µM tick and overlayed using ImageJ 3D Project. The movie shows only donor (blue) and ICE*Bs1* (red) to have an easier view of the transconjugants. Donor bacteria appear blue to purple and transconjugant appear bright red. The scale bar represents a size of 10 µm. The video shown is representative of conjugative clusters observed.

